# Mitotic chromosomes are mechanically connected by chromatin-based interchromosome linkers

**DOI:** 10.64898/2026.08.31.748267

**Authors:** Mingxuan Sun, Ron Biggs, Omar Akhtar, Gabriel Quintero Plancarte, Lisa L. Hua, John F. Marko

**Affiliations:** Department of Molecular Biosciences, Northwestern University, Evanston, Illinois 60208-3500; Department of Biology, Sonoma State University, Rohnert Park, CA 94928; Department of Physics and Astronomy, Northwestern University, Evanston, Illinois 60208-3500

## Abstract

The spatial organization of chromosomes is crucial for gene regulation and genome stability. Metaphase chromosomes appear physically discrete from one another in conventional chromosome preparations. However, interchromosomal connections, or “linkers”, have been observed among mitotic chromosomes, and may play important roles in coordinated chromosome movements and genome stability. We use micropipette-based isolation and manipulation to analyze interchromosome linkers between mammalian mitotic chromosomes. Pulling an isolated chromosome reveals that thin linkers connect them; CREST staining indicates their locations to be at centromeres. The linkers display linear elasticity and have a length-doubling force of approximately 300 pN, comparable to the length-doubling force of an entire metaphase chromosome. Enzymatic treatments demonstrate that the linkers are disrupted by DNase, but not by RNase, protease, or microtubule polymerization (spindle fiber) inhibitors, indicating that linker connectivity is based on DNA. Immunofluorescence experiments indicate that histones and topo II are on the linkers, with topo II organized into discrete foci spaced by approximately 0.4 Mbp. After prolonged metaphase arrest using spindle inhibitors isolated genomes do not display interchromosome linkers. Finally, experiments on intact cells show that interchromosome linkers containing CENP-B can be observed between mitotic chromosomes, indicating they are not an artifact of genome isolation.

**Significance:**

1. Chromosomes are connected by DNA-based linkers during mitosis
2. Interchromosome linkers are thin (roughly 100 nm), highly elastic, are attached near centromeres, and may serve to hold chromosomes together through mitosis
3. Interchromosome linkers contain histones and Topo II, the latter organized into discrete loci spaced by roughly 0.5 Mbp

**Highlight Summary (for Table of Contents):** This paper examines the composition, size and mechanical properties of interchromosome linkers, which are DNA-based filaments linking chromosomes together during mitosis. The linkers are elastic, contain approximately a few Mb of DNA, and may function to hold genomes in set conformations through cell division.

## Introduction

Mitotic chromosomes are often viewed as physically separated gene linkage units, partly due to their discrete morphology in conventional chromosome spreads. This perception is supported by the finding that chromosome-sized DNA molecules have been isolated from cells, indicating each chromosome is composed of one contiguous linear piece of DNA molecule in yeast and Drosophila (Kavenoff and Zimm, 1973; Carle and Olson, 1984; Schwartz and Cantor, 1984). However, physical connections or interchromosome “linkers” between mitotic chromosomes have been observed in a wide range of cell types (human, deer, bovine, chaffinch, newt, mouse) (Hoskins, 1968; Korf and Diacumakos, 1978; Maniotis *et al*., 1997; Saifitdinova *et al*., 2000; Poirier and Marko, 2002b; Kuznetsova *et al*., 2007; Sun *et al*., 2011; Sun *et al*., 2018; Potapova *et al*., 2019; Biggs *et al*., 2025).

In addition to interchromosome linkers between mitotic chromosomes of somatic cells, it has been shown that chromosomes are connected during meiotic metaphase of mouse, monkey, and plant cells (Klasterska *et al*., 1976, 1977; Klasterska, 1978; Hornick *et al*., 2015; Biggs *et al*., 2020; Liu *et al*., 2025). It is to be emphasized that we are concerned with pre-anaphase linkages between separate mitotic chromosomes, hence the term interchromosome “linkers”, and not the DNA-containing “anaphase bridges” that can occur between replicated chromatids (Finardi *et al*., 2020).

Spatial organization of chromosomes inside of the nucleus is crucial for nuclear function. It is known that interphase chromosomes occupy distinct chromosome territories in the nucleus, and that positioning of a chromatin locus affects its gene expression (Cremer and Cremer, 2001; Brickner and Walter, 2004; Bolzer *et al*., 2005). Interchromosome linkers may be related to fundamental questions: Are mitotic chromosomes randomly arranged in the cell, or do they have preferred neighbors? How is the spatial arrangement of chromosomes maintained during the cell cycle, especially during mitosis?

Conceivably, interchromosome linkers could play a role in preserving chromosome order through cell division. It is still an open question whether the positions of chromosomes relative to each other have any preference to their neighbors (Miller *et al*., 1963; Nagele *et al*., 1995), or are random (Korf and Diacumakos, 1977; Allison and Nestor, 1999; Dozortsev *et al*., 2000). Recent studies have shown that during mitosis, there is a preference to “antipair” or separate homologous chromosomes into distinct clusters with correlated motion of CENP-A foci (Hua and Mikawa, 2018; Hua *et al*., 2022; Cai *et al*., 2025). This antipairing may suppress illegitimate recombination of homologous chromosomes and suggests that there exist mechanisms such as interchromosome linkers that help to maintain supra-chromosomal genomic organization and genome stability.

An early hypothesis was that interchromosome linkers were composed of spindle fibers connecting different chromosomes (Hoskins, 1968). However, experiments showed the linkers to be DNase sensitive and colchicine resistant, suggesting that they were DNA-based fibers (Korf and Diacumakos, 1978; Maniotis *et al*., 1997). The width of interchromosome linkers has been estimated to be ≈100 nm (Hoskins, 1968; Marko, 2008). It is unclear whether interchromosome linkers share the same basic architecture as nucleosome-containing chromatin fibers found in mitotic chromosomes.

Interchromosome linkers are absent in most chromosome spreads, possibly because they break during cell lysis and centrifugation, resolve during metaphase stalls, or are simply too thin to observe using fluorescent dyes. It has been reported that hypotonic solutions facilitate rupture of interchromosome linkers (Klasterska, 1978).

The location of interchromosome linkers along chromosomes is controversial. Some studies support the hypothesis that interchromosome linkers run from one centromere to another (Hoskins, 1968; Korf and Diacumakos, 1978), while telomeric attachments were also reported (Jaffray and Geneix, 1974; Myhra and Brogger, 1975). The composition of the linkers has also been controversial. It has been reported that the linkers co-stain with α-satellite DNA binding protein, CENP-B, and RNA helicase p68 (Kuznetsova *et al*., 2007). Centromeric satellite DNA has been found in interchromosome linkers (Dozortsev *et al*., 2000; Saifitdinova *et al*., 2000; Saifitdinova *et al*., 2001; Kuznetsova *et al*., 2007).

A recent study used super-resolution imaging to examine interchromosome linkers, focusing on the hypothesis that some of them are based on ribosomal DNA (rDNA) (Potapova *et al*., 2019). rDNA-containing linkers between acrocentric human chromosomes were indeed observed, and the rDNA linkers are dependent on and contain the ribosomal gene transcription factor, UBF. These linkers were proposed to be a result of heavy entanglement of rDNA loci in nucleoli and are relevant to the five acrocentric human chromosomes (chromosomes 13, 14, 15, 21 and 22). However, linkers have been observed connecting the entire genome, and therefore, there must also be non-rDNA-containing interchromosome linkers associated with the remaining 18 non-acrocentric chromosomes.

In this study of interchromosome linkers, we used micropipette-based manipulation to extract mitotic genomes from human HeLa and mouse embryonic fibroblast (MEF) cells, using methods we have developed to isolate individual chromosomes (Poirier and Marko, 2002b; Sun *et al*., 2011; Sun *et al*., 2018; Biggs *et al*., 2025). Without drug treatments we always observe physical linkages between chromosomes, with the genomes organized into clusters or “bundles” of mitotic chromosomes. Although their mechanical effects are obvious, those linkers are barely visible under phase contrast, indicating a width in the range of approximately 100 to 200 nm. We find that the linkers are anchored at the centromere of each chromosome, and can be disrupted by DNase, but not RNase, protease, or colchicine treatments. We measured the mechanical response of the linkers, and we found a doubling force of ∼300 pN, comparable with the doubling force of whole chromosomes. Immunofluorescence staining of chromosome-associated proteins reveals that histones and topo II are associated with the linkers. Nocodazole and colchicine-treated cells do not have linkers present, indicating that they are cleaved or resolved during prolonged mitotic arrest. We also have done high-resolution fluorescence microscopy experiments imaging interchromosome linkers in intact cells, with the result that we observe CENP-B staining along the linkers between centromeric regions.

## Materials and Methods

### Cell culture

HeLa (human cervical adenocarcinoma cell, ATCC #70016359) and MEF (mouse embryonic fibroblast NIH /3T3, ATCC #CRL-1658) cells were cultured in DMEM with 10% FBS (Gibco), 100U/ml penicillin/streptomycin (Fisher Scientific) in a 37°C incubator supplemented with 5% CO_2_. Experiments were performed on ∼70% confluent samples.

### Microscopy for chromosome micromanipulation

Cells were transferred to and grown in sample dishes were prepared using #1 microscope glass onto which 25 mm-diameter rubber O-rings were affixed using paraffin. Pipette micromanipulation and imaging were done on the stage of an inverted microscope (IX70; Olympus) using a 60X 1.4 NA oil immersion objective at 30 °C with a temperature-controlled heater attached to the objective. Metaphase cells were identified by phase–contrast imaging. For the cases of mitotic arrest, cells were treated with 2.5ng/ul colchicine (Sigma-Aldrich) or 1ng/ul nocodazole (Sigma-Aldrich) overnight (16 hours).

HeLa cell imaging for micromanipulation was done using a CCD camera (Pelco, DSP B&W) with images acquired by a frame grabber (IMAQ PCI-1408, National Instruments); or a CCD camera (Ixon3, Andor). In brief, prometaphase cells were identified by phase-contrast imaging, and a micropipette filled with 0.05% v/v Triton X-100 (Fisher Scientific) in PBS pipette two micropipettes were positioned into the sample dish controlled by a motorized manipulator (MP-285; Sutter Instrument Co.) to destabilize the cell membrane by microspraying. After the chromosomes escaped from the cell, two micropipettes were positioned in the sample dish. One of the two pipettes was pulled with a shorter taper so it was stiff, and the other one was pulled with a long taper to have a softer tip with spring constant of 50 pN/µm. The stiff pipette was used to catch one end of a single chromosome using aspiration. Then the floppy pipette was attached to the other chromosome end, with a linker connecting them.

For force measurements, the stiff pipette was moved at a rate of ∼2 µm/min, slow enough to avoid viscoelastic effects and therefore to equilibrate forces through the chromosomes and linkers between the pipettes (Poirier *et al*., 2000; Poirier and Marko, 2002b, a). Bending of the force pipette was recorded to monitor the force applied on chromosomes and linkers. Each extension-relaxation measurement was repeated at least 3 times to ensure its reproducibility. Micromechanical data were collected using image analysis software written in Labview (National Instruments). Pipette stiffness calibrations were carried out using a reference pipette of approximately 300 pN/µm stiffness that was itself calibrated using an electronic nanonewton force sensor (FemtoTools, FT-S100) while observing pipette bending on a microscope.

### Biochemical reactions on isolated genomes carried out by microspraying

MNase and type II restriction enzymes were used to induce cuts in double-stranded DNA, using microspraying from a pipette loaded with approximately 30 μL of reagent in reaction buffer. MNase (Fisher Scientific) was prepared at 1 unit/µl in PBS with 1 mM CaCl2. The restriction enzymes used were AluI (AG↓CT, New England Biolabs), and PvuII (CAG↓CTG, New England Biolabs), all of which produce blunt DNA cuts. Restriction enzymes were prepared at a concentration of 1 units/μl in appropriate reaction buffers (Tris/HCl, pH 7.5; 50 mM NaCl; 5 mM MgCl_2_). Trypsin (Sigma-Aldrich) was prepared at a concentration of 200 nM in PBS. Proteinase K (Promega) was prepared at a concentration of 200 nM in PBS with 1 mM CaCl_2_.

Human topo IIα enzyme was purified as described (Kawamura *et al*., 2010) and was diluted in its activity buffer (10 mM Bis-Tris-propane/HCl, pH 7.9, 120 mM KCl, 2 mM MgCl_2_, 0.5 mM EDTA, and 30 ng/μl BSA) with 1 mM ATP (Thermo Fisher Scientific) to a final concentration of 90 ng/ul. Topo II decatenation activity was tested using a kDNA (Topogen) decatenation assay.

Colchicine (Sigma-Aldrich) was diluted in PBS to a final concentration of 5 ng/ul for direct microspray experiments. Colchcine and nocodazole were diluted in DMEM to a final concentration of 2.5 ng/ul for HeLa cell mitotic arrest. Triton X-100 (Sigma-Aldrich) was diluted in PBS to a final concentration of 0.05% (V/V).

### Immunofluorescence staining of isolated chromosomes and linkers by microspraying

Primary and secondary antibodies were diluted 1:100 in 50% PBS with 0.1% Casein. Antibody solutions were loaded into the spray pipettes, pulled and cut to have a ∼5 µm opening. Microspray pipettes were mounted on a three-axis manual manipulator (Taurus, World Precision) and positioned manually near the isolated chromosome. Microspraying of reagents was carried out for 10 mins unless otherwise specified with applied pressure of 10-100 Pa and stopped for several minutes to allow spray solution (total volume of ∼5 ul) to diffuse away into the 1.8 ml sample dish (Poirier and Marko, 2002c; Pope *et al*., 2006; Kawamura *et al*., 2010). After single chromosome extraction, chromosomes were sprayed with the blocking solution (PBS with 0.5% Casein), primary antibody and secondary antibody sequentially. DAPI (Invitrogen) was diluted to a final concentration of 50 ng/µl.

Primary antibodies used in microspraying experiments were: rabbit polyclonal SMC2/hCAP-E antibody (07-710, upstate), rabbit polyclonal Topo IIα antibody (TG2011-1, Topogen), mouse polyclonal histone antibody (MAB3422, Millipore). Secondary antibodies used were Alexa 488 donkey anti-mouse IgG (Invitrogen), Alexa 594 goat anti mouse IgG (Invitrogen), and Alexa 488 goat anti-rabbit IgG (Invitrogen). Centromeres were detected using Texas Red-conjugated CREST antibody (15-235-T, Antibodies Inc).

### Intact cell immunofluorescence staining

HeLa cells were grown on slides to 70-90% confluence. Slides were fixed with 4% paraformaldehyde (PFA) in 1x phosphate buffered saline (PBS) solution in a Coplin jar at room temperature (RT) for 10 min. Slides were then washed 3 times, each for 5 min with chilled 1x PBS. Following the 1x PBS washes, an antigen retrieval (AR) buffer (10mM Sodium Citrate, 0.05% Tween-20) was added to the slides and heated in a steamer for 10 min. After discarding the AR buffer, a permeabilization buffer (0.25% Triton X-100 in 1x PBS) was added for 10 min at RT. Three more 1x PBS washes at RT were completed. The slides were then placed in a moisture container and incubated in blocking buffer (10% goat serum, 0.1% Tween-20 in 1x PBS) for 30 min. Next, the blocking buffer was removed, and the slides were incubated with primary antibodies to human CENP-B (1:1000, Abcam ab25734), and γ-tubulin (1:1000, Abcam ab27074) in a 0.1% Tween-20, 10% goat serum, 1x PBS solution. Parafilm was used to cover the slides, and the slides were incubated at 4°C overnight.

The following day, the primary antibody was removed, and the slides were washed three times with 1x PBS at RT. The slides were then incubated with secondary antibodies to goat anti-rabbit IgG Alexa Fluor 488 (1:500, Abcam ab150077) and goat anti-mouse IgG Alexa Fluor 594 (1:500, Abcam ab150116) in a 0.1% Tween-20, 1% bovine serum albumin (BSA) and 1x PBS solution. Hoechst DNA counterstain was also included with the secondary antibody incubation. Slides were then covered with parafilm, and placed in the moisture container in the dark for 1 hr. Then, 3 more 1x PBS washes were completed. The slides were mounted with ProLong Gold antifade (Invitrogen, P36930) before addition of cover slips.

### Intact cell imaging and analysis

Mitotic cells were identified by the higher intensity of DNA staining in comparison to interphase cells. At prometaphase, chromosomes are arranged in a circular, or rosette structure (Magidson *et al*., 2011).

Fixed cells were observed and captured with a 63x oil immersion objective at 2.0x digital zoom on a confocal microscope (Leica TCS SPE, DM2500). A 1024x1024 frame size was used in obtaining z-stacks, and images were processed on Leica Application Suit X software (3.5.2.18963). Imaris image analysis software (Bitplane) was used for 3D reconstruction and measurement of CENP-B foci. Imaris deconvolution algorithm was applied, and all channels’ fluorescence levels were thresholded by including 90% of each signal. CENP-B foci were identified as 0.50 um of fluorescence signal/diameter as previously described (Cai *et al*., 2025).

### Statistics

For bar graphs and reported values we display and state the average ± standard error of the mean. The numbers of trials for each measurement are reported either in the figures, the figure captions, tables, or table captions. The set of linker lengths and stiffnesses determined in micromanipulation experiments are reported in the Supplemental Table. Statistical significance was determined by a Student t test, with significance defined as P < 0.05. Results of enzyme digestions were scored as 1 for cut or 0 for not cut (loss of mechanical connection observed); outcomes are provided in Table 1.

**Table 1.**
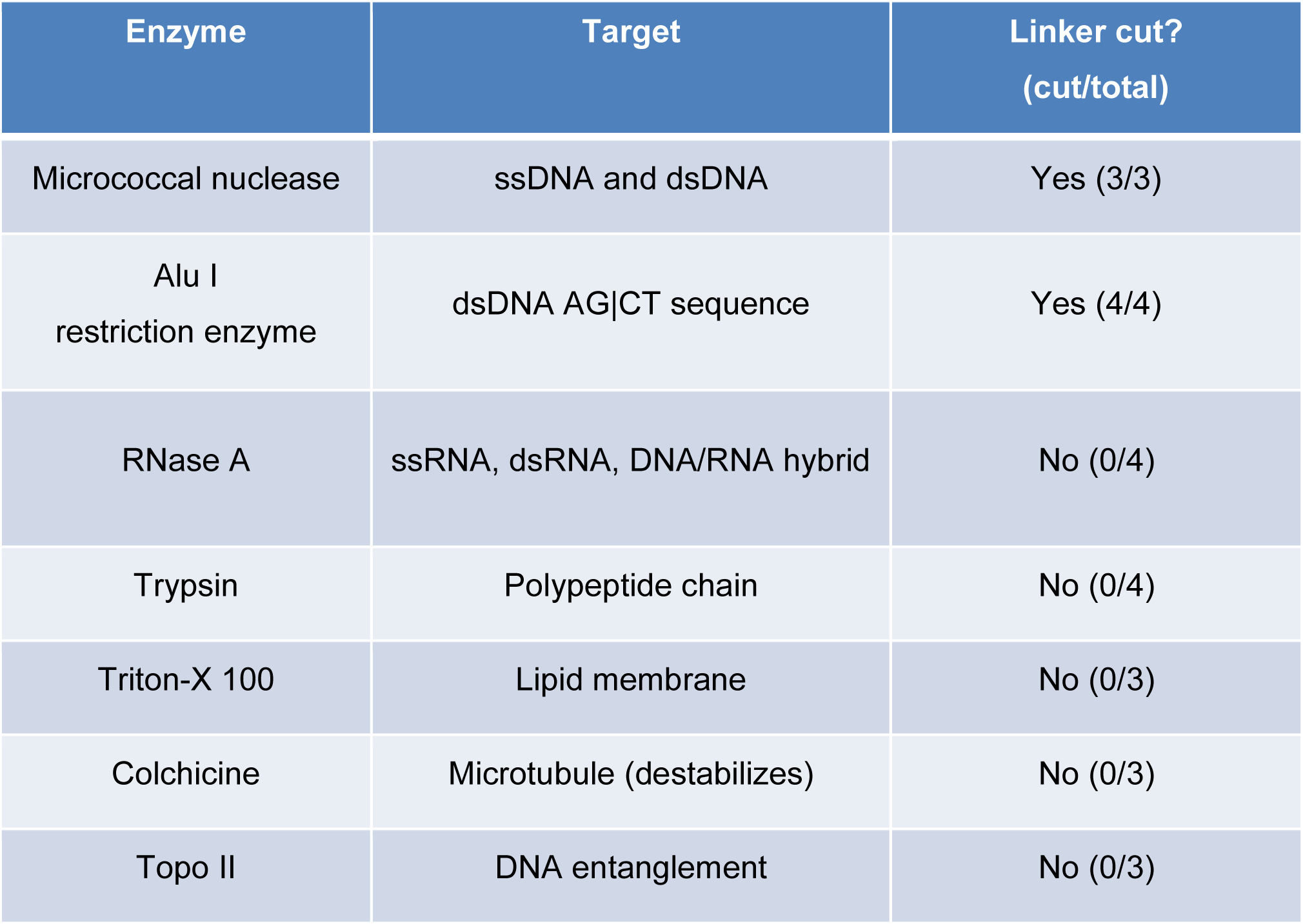
Enzyme digestion of interchromosomal linkers. Isolated linkers were subjected to a microspray of enzyme in activity buffer. Results are quantified in terms of cuts/attempted experiments.

In intact cell fluorescence experiments, binomial logistic regression was used to assess the effect of cell phase on the probability of the presence of linkers. A general linear model was used to assess whether the percentage of the number of linkers per CENP-B focus differed between cell phases. The linker per CENPB percentage data were log transformed to meet parametric assumptions. Statistical analysis was performed using statistical analysis software, JMP Pro (SAS Institute, Raleigh, NC).

## Results

### Interchromosome linkers physically connect centromeres of mitotic chromosomes

We used two glass micropipettes controlled by motorized manipulators (Sutter, MP-285) to extract chromosomes from living HeLa cells at prometaphase, as described previously (Poirier and Marko, 2002b; Sun *et al*., 2011; Biggs *et al*., 2025). In brief, a spray pipette filled with PBS with 0.05% Triton X-100 was used to gently open the cell membrane; individual chromosomes could then be caught by another pipette using aspiration (Fig. 1A). We have used this approach to isolate single chromosomes from mitotic cells, and to study the micromechanics of mitotic chromosomes (Fig. 1A’). Pulling on one chromosome away from a cell almost always resulted in sequential removal of all chromosomes, as if they were physically connected by thin threads. When a micropipette was attached to one of the chromosomes and was then moved out of the cell, neighboring chromosomes were pulled out from the cell sequentially (Fig. 1B, B’), indicating that they were physically connected.

**Figure 1.**
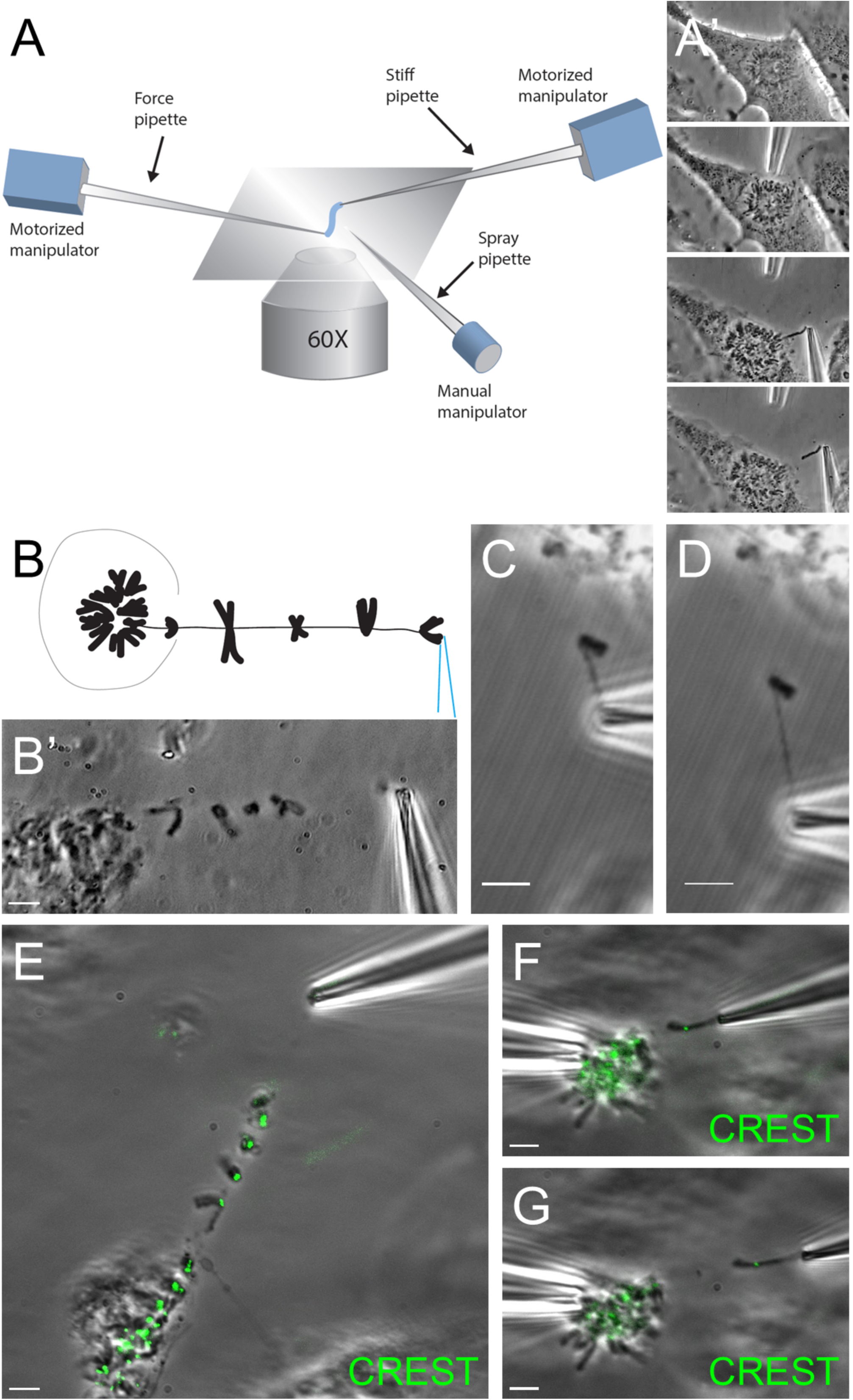
Physical linkages are present between human mitotic chromosomes. (A) Micropipette-based chromosome micromanipulation setup. Two micropipettes controlled by motorized manipulators are used to sequentially open the cell membrane and surgically remove mitotic chromosomes from the cell. Cells grow on a #1 microscope cover glass and are not synchronized. Individual prometaphase cells were identified by phase-contrast microscopy. Right panel shows phase contrast images of the sequential removal of a single mitotic chromosome from a prometaphase cell, with rosette-shaped structure of the connected chromosomes. Following extraction, one or more mitotic chromosomes can be suspended between two pipettes in the aqueous tissue culture medium in which the cells are grown. Bar = 5 μm. (B) Microsurgical removal of tandem chromosome arrays tethered at the centromeres. One chromosome is captured by a micropipette by aspiration, removing one chromosome from the cell results in the movement of all chromosomes in that cell. (C and D) Phase contrast images of a single chromosome captured by its end by a glass micropipette, before (C) and after (D) stretching. The isolated chromosome is kinked in the middle, due to the interchromosome linker. From C to D, only half of the chromosome arm is stretched by applied force. (E-G) Overlay images of phase-contrast (grey) and CREST immunostaining (green) on the chromosome cluster from one cell (E). A single chromosome captured by a glass micropipette before (F) and after stretching (G).

To investigate the location of the tethering point to a chromosome, we attached a micropipette to a free end of a chromosome and examined the connectivity of the linkers. One chromosome pulled away from the rest of the genome by a micropipette attached to one end showed in most cases the linkers were tethered to the middle of a chromosome, rather than to a chromosome end (Fig. 1C). More applied force resulted in stretching of the linker, as well as half of the chromosome arm between the linker and the micropipette, without affecting the rest of the chromosome arm (Fig. 1D). This type of experiment was repeated with the same results 4 times.

To verify that chromosomes were interconnected at the centromeres, a centromeric stain (CREST) was microsprayed onto the chromosomes (Fig. 1E-G, CREST staining in green). Overlaid images of phase contrast and CREST staining indicated that the interchromosome linkers ran from one centromere to another (Fig. 1E). In cases where a centromeric or telomeric attachment was not visually clear, mechanical pulling experiments after CREST staining showed that applied force on one end of the chromosome only led to the stretching of half of the chromosome, without affecting the other half (Fig. 1F-G). Our results demonstrate that chromosomes are linked from one centromere to another (repeated 5 times with same result).

### Linkers are sensitive to DNase digestion, but are RNase and colchicine resistant

We examined the nature of the linkers by directly spraying fluorescent dyes or reagents onto them using a spray pipette (Figs. 2-4). Although the linkers were barely visible under phase contrast (Fig. 2A), DAPI staining of a cluster of chromosomes clearly showed a thin thread connecting different chromosomes, indicating the presence of contiguous DNA filaments (Fig. 2B). We note the very strong overexposure that occurs for chromosomes due to the massive amount of DNA and DAPI fluorescence (microspraying leads to staining of the linkers and all nearby chromosomes), at the level of imaging exposure needed to observe interchromosome linkers, which makes them extremely difficult to observe in an intact cell where all the chromosomes are packed closely together. Our ability to stretch out the interchromosome linkers using micromanipulation is key to our ability to observe and analyze them.

**Figure 2.**
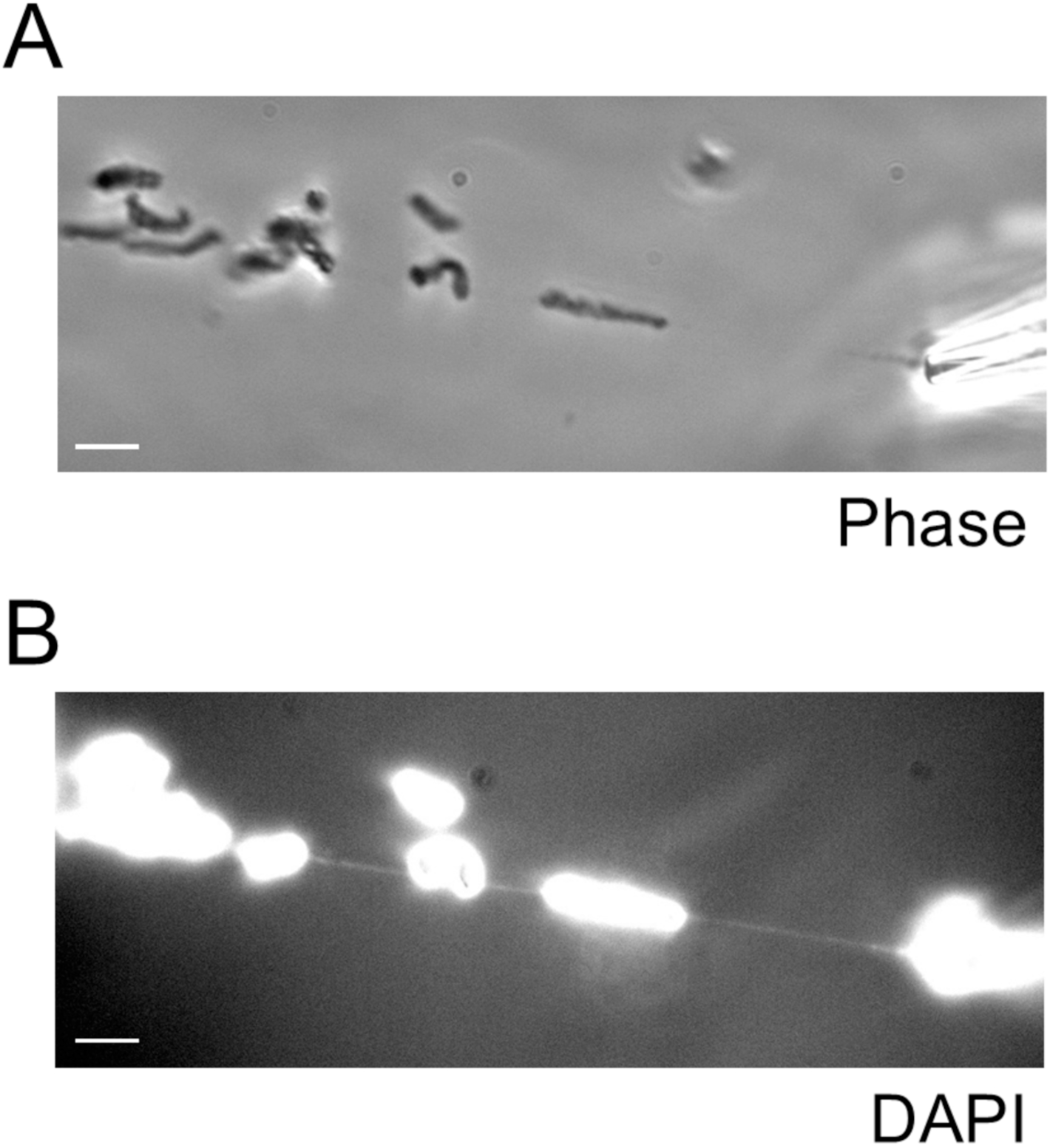
Metaphase chromosomes are connected by DNA linkers. Phase contrast (A) and DAPI staining (B) of the same chromosome cluster microsurgically removed from HeLa cells using micropipettes.

**Figure 3.**
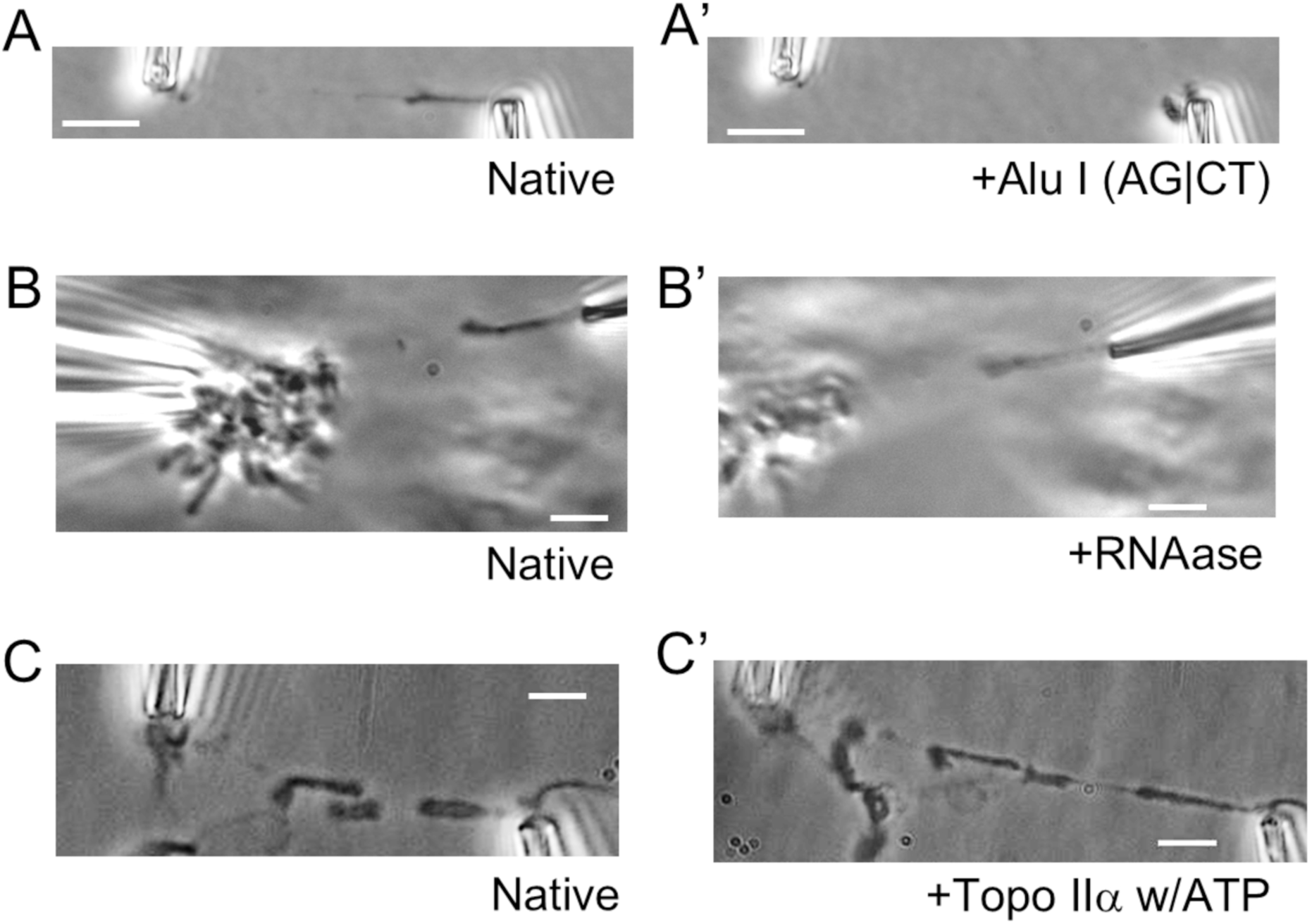
Interchromosome linkers are cut by DNase but not RNase. (A) DNA cutter Alu I cuts interchromosome linkers. Alu I is a blunt cutting restriction enzyme that cuts AG|CT sequence on DNA. Experiments with MNase led to similar results (data not shown). (B) RNase digestion does not have a significant effect on the linker. RNase A was diluted 1:100 and sprayed on to the chromosome linker for 20 mins. (C) Topo II + ATP spray on the linker. 90 ng/µl human Topo II with 1mM Mg·ATP was introduced onto the chromosome-linker cluster. Spray was stopped after 20 mins. Bar = 5µm.

**Figure 4.**
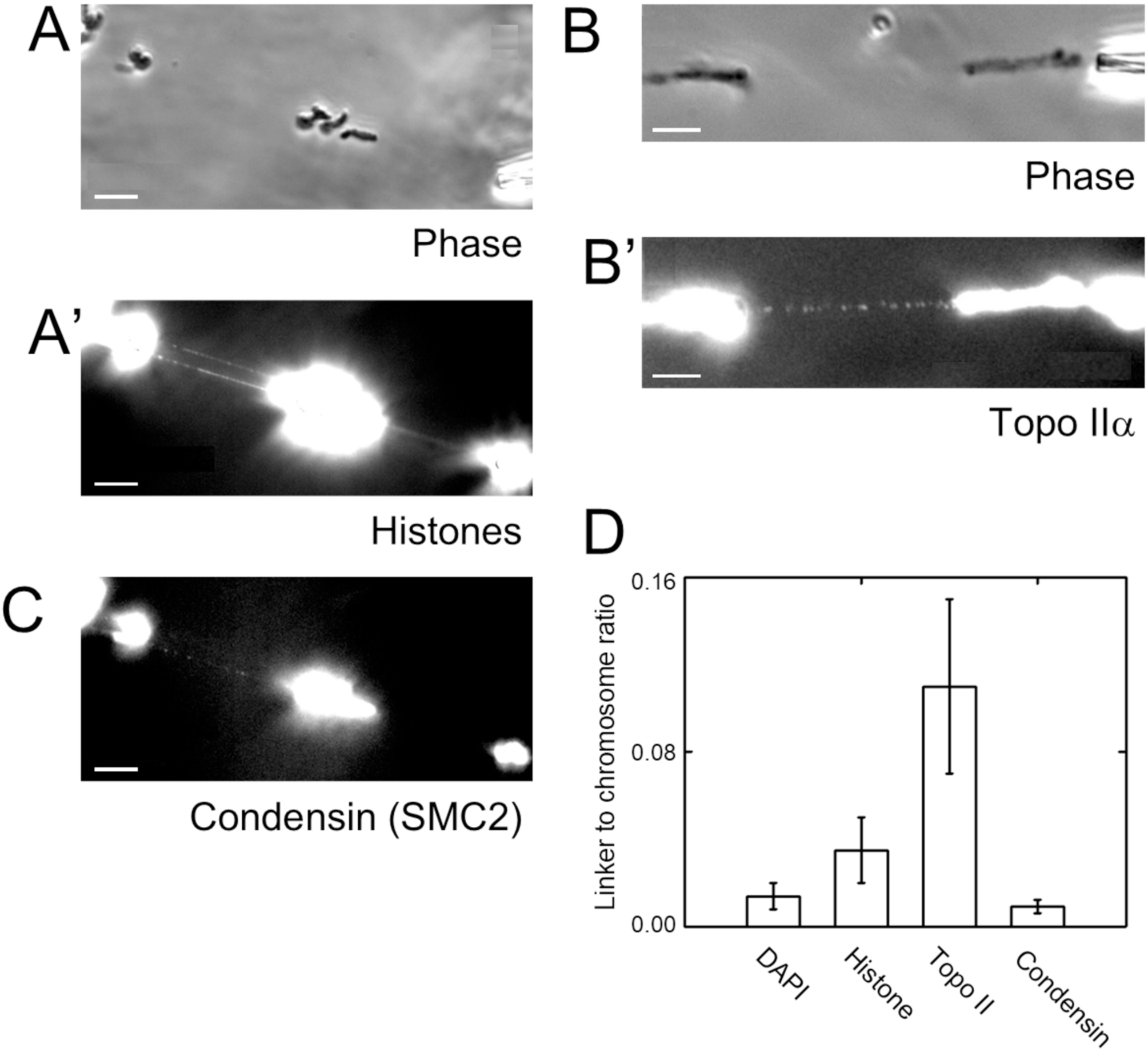
Immunofluorescence staining of interchromosome linkers. (A) Phase-contrast images of the chromosome cluster connected by interchromosome linkers; (A’) shows immunofluorescence staining of histones on same linker-chromosome structure. Bar = 5 µm. (B) Phase contrast image of two chromosomes connected by interchromosome linkers; (B’) shows immunofluorescence labeling of Topo IIα and formation of approximately 10 puncta along the extended linker. Bar = 5 µm. (C) Only low levels of immunofluorescence staining for SMC2 (condensin) are observed along interchromosome linkers. (D) Ratio of total fluorescence for linkers and an adjacent mitotic chromosome for DAPI, histone, TopoIIα and SMC2 (condensin) (see also Table 2).

**Table 2.** Immunofluorescence labeling of interchromosomal linkers. Each row shows integrated fluorescence counts along a linker (arbitrary units, after background subtraction) and ratio to integrated fluorescence for an adjacent chromosome), averaged over a series of trials. For PICH and cohesin the measured fluorescence value is within standard error of zero, indicating no difference between the linker fluorescence and the background, with a zero value of ratio to chromosome fluorescence.

| IF target | Linker fluorescence<br>(arbitrary units) | Linker to adjacent<br>chromosome ratio | N |
| --- | --- | --- | --- |
| PICH | -15 ± 26 | 0 | 4 |
| cohesin | 2.2 ± 7.6 | 0 | 4 |
| condensin | 77 ± 46 | 0.0093 ± 0.0063 | 6 |
| Topo II | 265 ± 63 | 0.11 ± 0.04 | 6 |
| DNA (DAPI) | 230 ± 34 | 0.014 ± 0.006 | 4 |
| histone | 1500 ± 480 | 0.035 ± 0.015 | 4 |

To test whether the linkers depend on DNA and not on proteins, e.g., spindle fibers (Hoskins, 1968), micrococcal nuclease (MNase, Thermo Scientific) or a Type II restriction enzyme,Alu I (AG|CT, Promega) were used to introduce double-stranded DNA breaks (Fig. 3A, A’). Before the enzyme sprays, the linker was stretched between two chromosomes under tension, as indicated by stretching of the chromosomes (Fig. 3A). After the Alu I spray, the linker was cut (Fig. 3A’). As the two chromosomes on each end of the linker retracted, either of the two pipettes moved away without any resistance (Fig. 3A’, experiment repeated 4 times with same result; MNase gave similar results for 3 trials, data not shown).

Similar experiments were performed using RNase A (New England Biolabs) (Fig. 3B, B’) which cuts single-stranded RNA, double-stranded RNA, as well as DNA/RNA hybrids. After 20 min of RNase spray on the stretched linker, indicated by the slightly stretched chromosome and the tension on the force pipette, the linker was still mechanically attached (Fig. 3B’). This experiment was repeated 4 times with the same result.

We also examined the possibility that chromosomes are held together by DNA entanglements, by directly spraying them with Topo IIα + ATP (Fig. 3C, C’). No significant change was observed on the linker after 20 min incubation (Fig. 3C’). This experiment was repeated three times with the same result.

We examined whether interchromosome linkers were dependent on other types of molecules, such as proteins (trypsin or proteinase K, no change in 4 experiments), spindle fibers using colchicine (no change in 3 experiments), and phospholipids of membranes using Triton X-100 (no change in 3 experiments). Using our spray method, we observed that none of those reagents had any noticeable effect on physical connectivity (Table 1).

### The linkers contain histones and Topo II, but little condensin

Interchromosome linkers are barely visible in phase-contrast images (Figs. 1-4), indicating an estimated width of ∼100 nm. Electron microscopy studies show an estimated linker width of 60 nm (Hoskins, 1968). To understand the organization of the linkers, we tested whether major chromatin organizing proteins such as histones, Topo IIα, and condensin were present in them (Fig. 4A-B’). A chromosome cluster was pulled away from the rest of the genome, with interchromosome linkers present in between the separate chromosomes (Fig. 4A), and then sprayed with antibodies to specific proteins. Linkers and nearby chromosomes were sprayed sequentially with primary antibodies to mouse anti-histone (Millipore, MAB3422), and rabbit anti-SMC2 (condensin) (Millipore 07-710) and then secondary antibodies (anti-rabbit IgG Alexa 488, anti-mouse IgG Alexa 564, Invitrogen) (Fig. 4A-A”).

We observed a significant amount of histone on the extended linkers with a varying distribution of fluorescence (Fig. 4A’). By comparison, there was little, if any, condensin fluorescence on the linker (Fig. 4A”). We observed a distinct periodic pattern of Topo II on the linker (Fig. 4B,B’). We examined the interchromosome linkers in mouse fibroblast (MEF) cells and found the same punctate pattern there (Fig. S1). We note that interchromosome linkers were stretched to be 3 to 5 fold longer than their native lengths to avoid the overwhelming fluorescence contribution from nearby chromosomes (Fig. 4A-B’).

We also tested the presence of several other chromosome-associating proteins including cohesin, and Plk 1-interacting checkpoint helicase (PICH), a protein which decorates anaphase bridges (Ke *et al*., 2011; Biebricher *et al*., 2013). Cohesin and PICH were not detectable on interchromosome linkers (Table 2).

To compare abundances of detected proteins to those on chromosomes, we measured total fluorescence of linkers and single adjacent chromosomes in the same image and then calculated the ratio of linker fluorescence to that of an adjacent chromosome, for DAPI, histone, Topo IIα, and condensin labeling (Fig. 4C, Table 2). Notably, the ratio of linker to chromosome fluorescence for Topo IIα was much higher for than condensin (Fig. 4C) suggesting Topo IIα to be enriched in linkers, and likely a key component organizing them.

### Interchromosome linkers have elastic stiffness similar to a whole mitotic chromosomes

To investigate the micromechanics of the linkers, we measured their force-extension responses. We used a force-sensing pipette to measure the force extension response of the linker. As it was difficult to grab the two ends of the linker by a micropipette, we attached two micropipettes to each of the two neighboring chromosomes, respectively, and used the resulting “chromosome handles” to stretch the linker (Fig. 5A). The force applied to the linker is equal to the force applied on the chromosomes, allowing measurement of the spring constant of the linker (or the chromosomes) by measuring linker extension and force-measuring pipette bending, and later calibrating the force-measuring pipette (Sun *et al*., 2011; Biggs *et al*., 2025).

**Figure 5.**
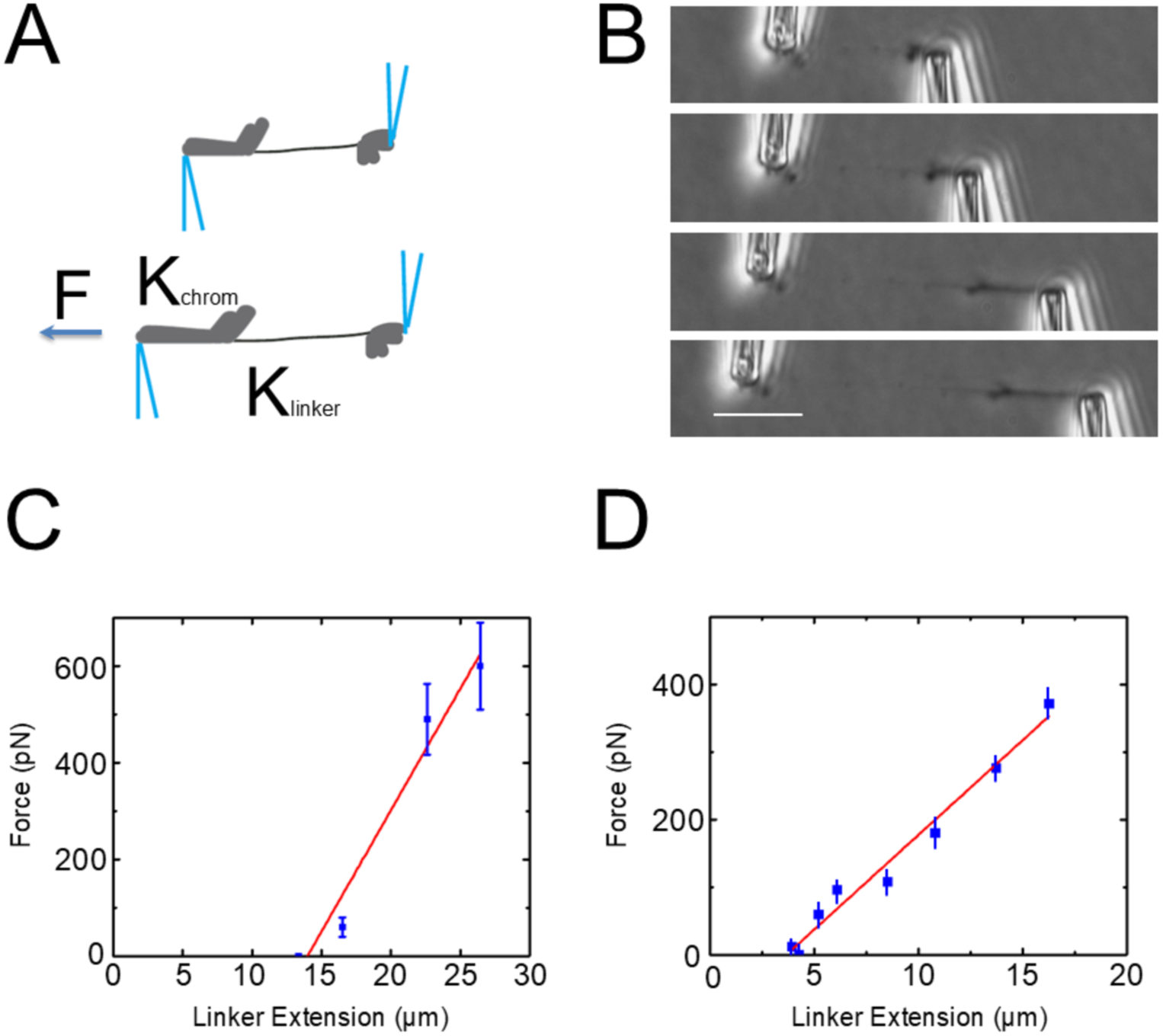
Micromechanics of the interchromosome linkers. (A) Schematic representation of the stretching experiment for the chromosome-linker-chromosome bundle. Under applied force F, the chromosome and linker have two different spring constants: K_chrom_ and K_linker_. (B) Phase contrast images of the stretching experiment for the chromosome-linker-chromosome bundle. (C) Force-extension curve of the linker shown in (B) with initial length of 14 µm. Red line shows the linear fit of the stretching curve, which gives a spring constant of 49 ± 7 pN/µm and a doubling force of 650 ± 100 pN. (D) Data for another trial on a separate linker of initial length 4 µm and spring constant 32 ± 3 pN/µm, indicating doubling force of 130 ± 10 pN.

In an example experiment two neighboring chromosomes held by two micropipettes showed a linker in between them (Fig. 5B). Stretching experiments showed that applying force to the combination of linker and chromosomes resulted in the stretching of the linker, as well as parts of the chromosome arms, indicating that the elasticity of the linker was comparable to that of the whole chromosomes. The linker was tethered at the centromere of each chromosome, resulting in only the part of the chromosome arm between centromere and micropipette being stretched under applied force as can be seen in Figs 1 and 5B. The force-extension behavior (Fig. 5C-D) is approximately linear over the two- to three-fold extensions that we studied and can be quantified in terms of the slope obtained from a linear fit as shown in Fig. 5C. The spring constant of this linker is just the fit slope of the linear fit, which is 49 ± 7 pN/µm.

The force at which this linker would be doubled in length if the initial linear elastic response were extrapolated, or its “doubling force” is a length-independent way to express linker extensibility and is related to the effective Young’s elastic modulus. Based on the linear fit, the doubling force for the data of Fig. 5C is 650 ± 90 pN. Based on N=9 trials we found the average doubling force to be 270 ± 60 pN. This is approximately 50 times higher than the 5 pN doubling force measured for a single chromatin fiber (Cui and Bustamante, 2000) (Fig. 5D), indicating that the linker is consistent with being a well-folded chromatin structure roughly 7 times thicker than a single nucleosome (i.e., about 70 nm thick) (Supplementary Table 1). It is also approximately the force constant observed for whole HeLa chromosomes in prior experiments (Sun *et al*., 2011).

### Colchicine-treated cells stalled at metaphase lack interchromosome linkers

We tested the possibility whether linkers were dependent on spindle fibers (microtubules) by directly exposing linkers to the microtubule polymerization inhibitor colchicine. We carried out removal of chromosome bundles connected by linkers from HeLa cells (Fig. 6A) and microsprayed colchicine onto them. The linkers were not disrupted by colchicine treatment, indicating the linkers were not supported by microtubule-based spindle elements (Table 1, 3 trials). This result is also consistent with our finding that the linkers are highly elastic, while microtubule filaments are relatively rigid and inextensible (Gittes *et al*., 1993).

**Figure 6.**
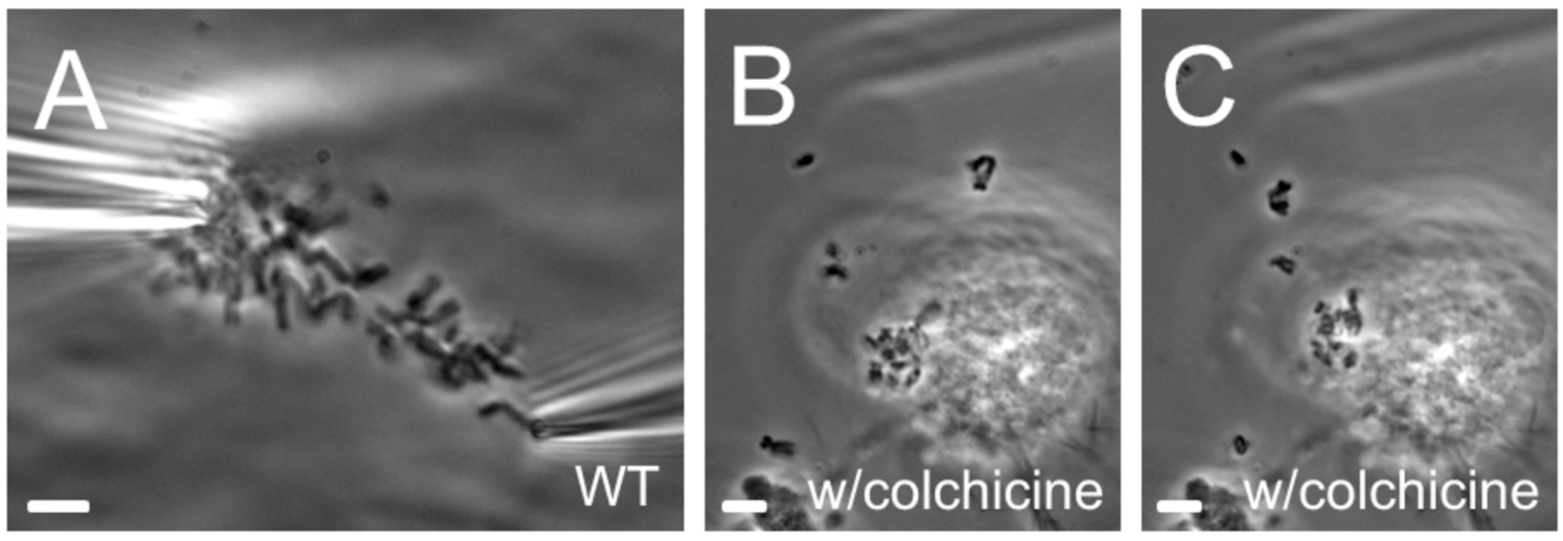
Spindle inhibitor-stalled cells lack linkers. (A) WT genome suspended between two micropipettes. Pulling one chromosome away from the rest of the genome results in the rearrangement of the whole genome. (B and C) Cells treated with colchicine for 14 hours. After the cell is opened, chromosomes diffuse out of the cell and are scattered in the culture media.

However, we did find that interchromosome linkers were much less common in cells treated with spindle inhibitors for >8 hours. We grew HeLa cells in DMEM with 1 ng/ul colchicine or 2.5 ng/ul nocodazole for 16 hours before opening mitotic cells (Fig. 6B-C). Following disruption of the cell membrane, we observed that many of the individual chromosomes escaped from the cells, freely diffusing in the culture medium without any apparent constraint between them (Fig. 6B-C).

### CENP-B-labeled interchromosome linkers can be observed in intact cells

The above experiments cannot address the possibility that interchromosome linkers somehow form during genome extraction and are an artifact of the isolation process. We therefore sought to visualize the linkers in intact cells. This is challenging since unlike the case of isolated genomes (Fig. 1-6), the chromosomes inside a mitotic cell are crowded together and the interchromosome linkers are unstretched; any dye that binds chromatin will generally overwhelm any imaging modality due to the massive amount of chromatin in a chromosome.

Given that prior experiments observe the pericentromeric DNA-binding protein, CENP-B, along interchromosome linkers (Kuznetsova *et al*., 2007), plus our observation that linkers emanate from the centromeric regions (Fig. 1E), we reasoned that probing for CENP-B might provide a signal along the interchromosome linkers that will not be overwhelmed by a general DNA or histone stain. Our strategy was to look for evidence of chromatin *between* chromosomal CENP-B foci along different chromosomes by combining information from DNA and CENP-B (centromere-pericentromere) staining.

Immunofluorescence (IF) staining of CENP-B protein was performed in HeLa cells, in conjunction with IF staining of γ-tubulin (centrosome) for identification of mitotic stage, and DNA (Hoechst) staining (Fig. 7). At prophase, CENP-B foci were observed as expected, and found to co-localize with bright (dense) Hoeschst-positive regions approximately 0.5 µm in diameter (Fig. 7A-A”’). These condensed Hoeschst-positive foci are expected due to the high compaction and density of DNA in centromeric heterochromatin (Biggs *et al*., 2025).

**Figure 7.**
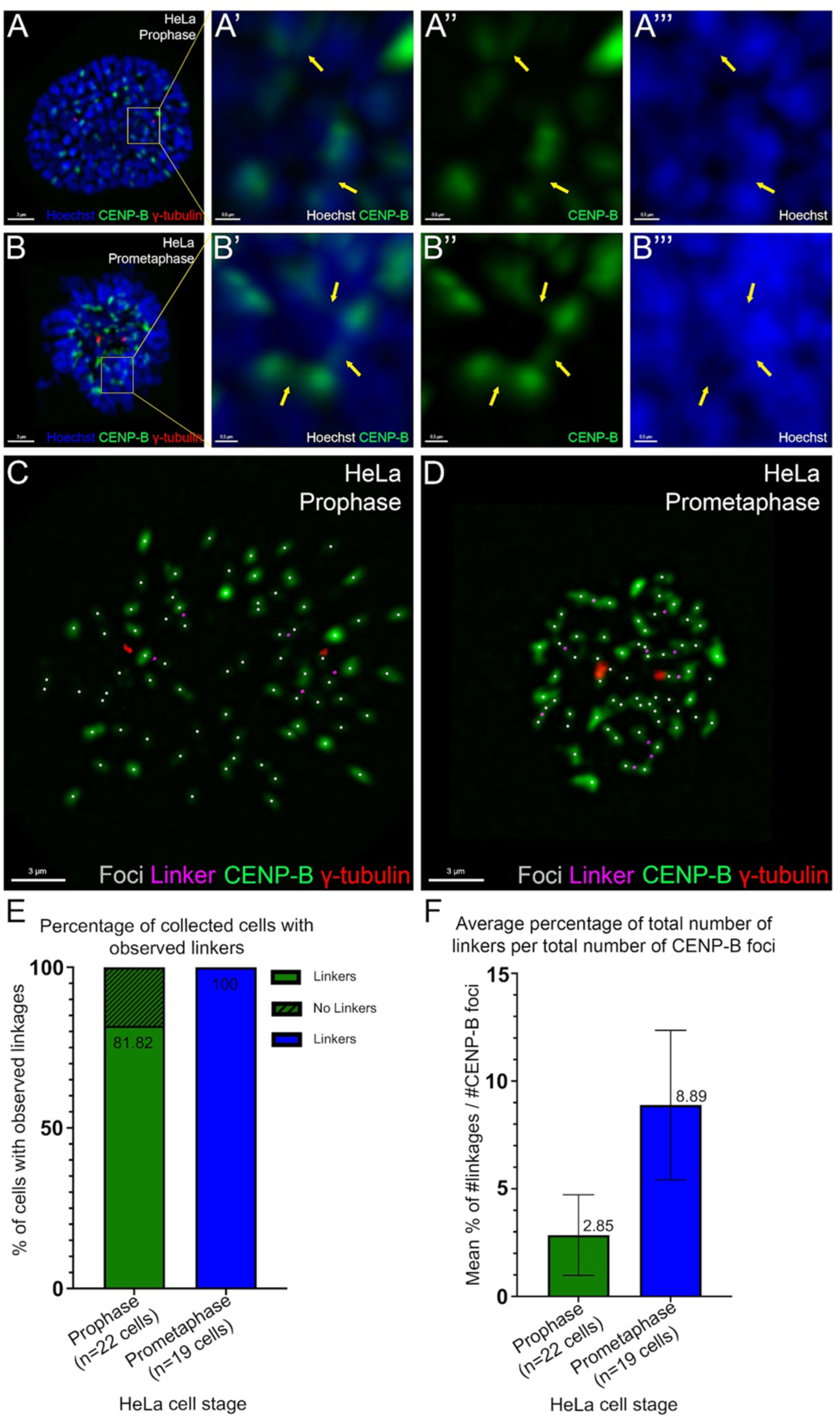
CENP-B stained interchromosome linkages in HeLa cells at prophase and prometaphase. **(A)** HeLa cell at prophase with immunofluorescence staining for CENP-B (green), *γ*-tubulin (red), and DNA counterstain (Hoechst, blue) (n=22 cells). Scale bar: 3 µm. **(A’)** As in (A), but of immunofluorescence staining for CENP-B (green) and DNA counterstain (Hoechst, blue). Yellow arrows indicate selected observed linkers. Scale bar: 0.5 µm. **(A’’)** As in (A’), but immunofluorescence staining for CENP-B (green). **(A’’’)** As in (A’), but DNA counterstain (Hoechst, blue). **(A’-A’’’)** Scale bars: 0.5 µm. **(B-B’’’)** As in (A-A’’’), but at prometaphase (n=19 cells). (**C**) Prophase HeLa nucleus of (A), with CENP-B (green), *γ*-tubulin (red) and interchromosome linkers (magenta) identified from CENP-B staining between chromosomes as in (A’-A’’’). White dots indicate CENP-B loci (pericentromeres). Hoechst/DNA staining not shown for clarity. **(D)** Prometaphase HeLa nucleus of (B); colors and markings as in (C). **(E)** Average percentage of collected cells at prophase and prometaphase with at least 1 observed linker. Linkers were observed in 81.82% (18/22) of cells at prophase and 100% (19/19) of cells at prometaphase. Linkers were not observed in 18.18% (4/22) of cells at prophase. **(F)** Average percentage of total number of linkers per total number of CENP-B foci at prophase (mean = 2.85, SD = 1.87) and prometaphase (mean = 8.89, SD = 3.47).

Between many of the CENP-B foci of different chromosomes and between tracks of the main DNA staining along chromosomes, diffraction-limited-thickness linear linkers of CENP-B positive staining were observed (Fig. 7A”’). We note that the single CENP-B foci resolved in imaging on intact cells correspond to *pairs* of sister chromatid pericentromeres, which are close together and appear as a single focus.

Similar observations were made for prometaphase cells (Fig. 7B-B’’’) with the same result that CENP-B-stained linkers could be observed between CENP-B foci of different chromosomes. These observations indicate that interchromosome linkers are identifiable and present between CENP-B foci in intact HeLa cells before (prophase) and after (prometaphase) nuclear envelope breakdown. Similar observations are impossible at metaphase due to the chromosomes being too close to one another on the metaphase plate to permit imaging of linkers.

We proceeded to quantify the linkers for prophase and prometaphase (Fig. 7A-B). The spatial resolution limit of the confocal microscope is 360 nm in xy (focal plane), so CENP-B stained linkers were only considered detected if they were at least this minimum xy thickness. CENP-B positive linkers were quantified by observing the top (xy) and side (yz, xz) views between at least two CENP-B foci (Fig. 7A). CENP-B-positive linkers that did not exceed the resolution width and did not run down the entirety of the stacked optical images of the CENP-B foci, were quantified as linkers (Fig. 7A’, A’’, B’, B’’). CENP-B stained regions that did not meet these criteria were not counted as linkers. Examples of linker-annotated prophase cells and prometaphase genomes are shown in Fig. 7C and 7D, respectively.

CENP-B positive staining of linkers were found between two or more CENP-B foci in 81.82% of Hela cells at prophase (n=18/22 cells) (Fig. 7E); 18.18% of cells did not show identifiable linkers (n=4/22 cells) (Fig. 7E; *X^2^* (1, n=41)=5.35, *P*=0.021). The average number of linkers per CENP-B focus was 3±2% (Fig. 7F; F_1, 39_=40.1, p<0.0001). CENP-B stained linkers also stained positive for DNA (Fig. 7A’’’). Observable CENP-B stained linkers between CENP-B foci were present in > 50% of prophase HeLa cells. Limitations in xy resolution combined with the crowding together of chromosomes in the prophase cell may constrain the ability to observe CENP-B linkers in all cells.

At prometaphase, all HeLa cells displayed at least one CENP-B positive linker between two CENP-B foci (n=19/19 cells) (Fig. 7E), with an average number of linkers per CENP-B focus of 9±4% (Fig. 7F). Both the incidence of observed linkers per cell and per CENP-B foci were larger for prometaphase than for prophase. This may indicate that linkers are more readily detectable with our imaging system at prometaphase as compared to prophase. Alternatively this may be a result of the rather orderly ring-like arrangement of chromosomes at prometaphase, positioning the CENP-B foci so that the linkers are more visible (Magidson *et al*., 2011). These observations of CENP-B stained interchromosome linkers in intact cells between CENP-B foci at prophase and prometaphase suggest that interchromosome linkers observed in isolated genomes are not the result of experimental *ex vivo* manipulations.

## Discussion

### Interchromosome linkers are structurally connected by DNA elements

The above experiments demonstrate that DNA, but not RNA, structurally connects different chromosomes in a metaphase cell (Fig. 2, Table 1). These DNA-based interchromosome linkers are decorated by chromosome associated proteins, including histones and Topo II (Fig. 4). Linkers are not sensitive to microtubule polymerization inhibitor exposure, indicating no dependence on microtubules. However, prolonged metaphase arrest (16 h incubation with microtubule polymerization inhibitors) leads to resolution of the linkers.

A mitotic chromosome is thought to contain one continuous piece of DNA, with its ends located in the telomeric end regions of the chromosome. The fact that the DNA-based interchromosome linkers are tethered near the centromeres of each chromosome suggests the hypothesis that the linkers are DNA loops connected by DNA entanglements, or possibly homologous recombination intermediates of the highly repetitive DNA found in centromeres or pericentromeres. We found that Topo IIα+ATP did not resolve the connections between the linkers indicating if interchromosome entanglements are the basis of the linkers, they are not easily resolved. However, it is still possible that such DNA entanglements are protected by other proteins, or are linked by recombination machinery, making them insensitive to Topo IIα+ATP. The linkers are also relatively thick, being barely visible in phase contrast, and may have appreciable chromatin folding that is not easily released by a reaction with Topo IIα + ATP.

The existence of interconnection between chromosomes may help to maintain an intact genome by linking adjacent chromosomes to make sure that all chromosomes are incorporated into the prometaphase rosette and participate in congression movements towards the metaphase plate during mitosis (Nagele *et al*., 1995; Maniotis *et al*., 1997). The DNA-based linkers connecting chromosomes may result in the formation of a network which uses DNA as tension transducing elements by directing coordinated chromosome movements to ensure faithful chromosome segregation (Maniotis *et al*., 1997). This elastic chromatin network may also help with unified movement of chromosomes to the spindle pole (Ingber *et al*., 1994). Moreover, interconnection between adjacent chromosomes may help to maintain the spatial order of chromosomes inside the cell (Nagele *et al*., 1995), and explain how movement of one chromosome can control the timing and movement of all chromosomes during mitosis (Alexander and Rieder, 1991; Li and Nicklas, 1995; Maniotis *et al*., 1997). When one chromosome is not properly attached to the spindle, the interconnections may allow it to be segregated due to its neighboring chromosomes (Nagele *et al*., 1995).

Notably, micromechanical study of the linker shows its force constant, the force needed to extend an object to double its length, to be comparable to those of human mitotic chromosomes (Sun *et al*., 2011; Biggs *et al*., 2025). Although the linker is approximately ten-fold thinner in diameter than human mitotic chromosomes (meaning roughly 100 times less cross-sectional area), linkers and chromosomes stretch by a comparable amount under the same external force. This provides a physical basis for the coordinated movement of chromosomes during mitosis: if the linkers were much softer than mitotic chromosomes, pulling on one chromosome would not lead to force transduction across the entire genome. At the same time, this indicates that the effective Young’s modulus of interchromosome linkers is much larger than the 500 Pa we have observed for mitotic chromosomes (Sun *et al*., 2011; Sun *et al*., 2018; Biggs *et al*., 2025), and is approximately 50 kPa. The linker Young’s modulus can also be estimated directly from the average force constant as Young’s modulus = (force constant)/(cross sectional area) = 300 pN / 10,000 nm = 30 kPa.

### Interchromosomal linkers are decorated by Topo II**α** clusters spaced by approximately 0.5 Mbp of DNA

We detected histones along interchromosome linkers, but only low levels of condensin (SMC2 staining, Fig. 4 and Table 2). We did detect a high level of Topo IIα, which was organized into remarkably evenly spaced puncta (Fig. 4B’, Fig. S1). Topo IIα enrichment on the linkers is consistent with the hypothesis that the interchromosome linkers are DNA loops held by DNA entanglements, and Topo IIα clusters may help with the regulation or resolution of the linkers at different cell cycle stages. Alternately Topo IIα may be playing a *structural* role, via its ability to directly mediate DNA compaction (Maniotis *et al*., 1997).

We can approximately estimate the approximate size of the linkers and the genomic distance between the Topo IIα clusters using the ratio of DAPI fluorescence (Table 2) on a linker relative to an adjacent whole chromosome (0.014±0.006). Taking the typical DNA content of a human or mouse replicated chromosome of 300±100 Mbp (two human chr 1 chromatids contain ≍500 Mbp; two of the small chr 22 chromatids contain ≍100 Mbp), this ratio suggests that an average linker contains approximately 5±2 Mbp of DNA. A numerically similar estimate results from using the histone fluorescence ratio (0.035±0.015, giving 10±3 Mbp); we average these results to estimate a mean linker content of 8±2 Mbp.

Given that we observed 10±2 Topo IIα clusters along linker (Fig. 4B’) we estimate that there is a cluster for each 0.8±0.2 Mbp. Assuming two chromatids in each linker, the clusters are spaced along each chromatid by approximately 0.4±0.1 Mbp. This is a DNA scale coincident with the 0.4 Mbp inferred from Hi-C data to be in large “outer” metaphase chromosomal loops (Gibcus *et al*., 2018). This 0.4 Mbp TopoIIα cluster scale is appreciably smaller than the roughly 5 Mbp scale for “condensin centers” observed along stretched human chromosomes using SMC2 immunofluorescence labeling (Sun *et al*., 2018); the condensin cluster scale is plausibly coincident with the 10 Mbp helical folding scale observed in Hi-C experiments (Gibcus *et al*., 2018). Given that the linkers contain Topo IIα clusters at a roughly 0.4 Mbp scale with little to no condensin (SMC2) and no large-scale folding, interchromosome linkers may represent an early stage of mitotic chromosome folding mediated mainly by Topo IIα, and independent of condensin.

It is worth comparing the approximate DNA we estimate in interchromosome linkers, 4±1 Mbp per chromatid, which corresponds to a total DNA length of 1200±300 μm, to the typical unstretched linker length which is 5 μm (average of unstretched lengths in Table S1). The linkers therefore correspond to a compaction level of roughly 1200/5 = 240-fold. While a large compaction consistent with the wide range of elasticity we have observed for linkers, this is well below the nearly 10,000-fold compaction for an entire mitotic chromosome, consistent with the linkers representing an intermediate level of chromosome folding.

### Spindle inhibitor-stalled cells lack interchromosome linkers

Direct exposure to colchicine does not disrupt the linkers, indicating that they do not directly depend on microtubules. However, the linkers are absent in cells which are stalled in metaphase by colchicine or nocodazole for an appreciable amount of time (16 hours). This finding could explain part of the controversy whereby several groups validate the existence of the linkers (Hoskins, 1968; Korf and Diacumakos, 1978; Maniotis *et al*., 1997) while others conclude that observation of linkers is an artifact of chromosome extraction, because chromosomes are not interconnected in colchicine-treated cells (Korf and Diacumakos, 1980). Our observation also helps to explain why those linkers are not often seen in conventional chromosome isolation methods, which generally utilize metaphase synchronization of cells using microtubule polymerization inhibitors.

In a prior work we observed that cells stalled in mitosis using colchicine develop highly compacted, condensin-overloaded, and well-resolved chromosomes (Sun *et al*., 2018). We hypothesize that topological resolution of inter-chromosome entanglements is coupled to condensin level on chromosomes: the more condensin on a chromosome, the more complete is the entanglement resolution process. Therefore, mitotic-stalled cells may have fewer or no interchromosomal linkers. The interchromosome linkers may be chromatin regions which are particularly resistant to entanglement resolution, but which eventually disentangle given a large enough amount of chromosomal condensin.

Importantly, it has been found that prolonged metaphase can induce DNA damage, possibly by activation of CAD DNase, which introduces random DNA cuts into the whole genome (Ganem and Pellman, 2012; Orth *et al*., 2012). Compared to the folded DNA in mitotic chromosomes, interchromosome linkers are more exposed and therefore more accessible to DNase, which may make them a prominent target for DNA cleavage during prolonged metaphase arrest. All these factors emphasize that basic aspects of chromosome organization can be changed by conventional mitotic arrest methods.

### Interchromosome linkers may be maintained through anaphase

Are interchromosome linkers permanent? How long do they last? When are they cleaved/resolved? Linkers between anaphase chromosomes have been observed in anaphase I of mouse spermatocytes (Klasterska, 1978), suggesting linkers may persist through the entire cell cycle. Interchromosome linkers during mitosis as analyzed here, have been suggested to arise from interphase interchromosome linkers visualized in microdissection experiments (Maniotis *et al*., 1997).

We have attempted to determine whether interchromosome linkers are present during HeLa cell anaphase. A phase contrast image of an extracted genome from an anaphase cell indicated that chromosomes continue to be connected by linkers (Fig. S2). This observation is crude and far less precise than our experiments at metaphase because of the confounding presence of mechanically strong cytoskeletal elements in anaphase (principally microtubules and actin), but Fig. S2 supports the case of (Klasterska, 1978) that interchromosome linkers persist through cell division.

### CENP-B-containing intercentromeric linkers are observable in intact cells

In confocal fluorescence microscopy experiments, we observed that CENP-B containing linkers can be observed between pericentromeric/centromeric regions of chromosomes (Fig. 7), in accord with metaphase spread visualizations (Kuznetsova *et al*., 2007). Our observation supports the hypothesis that interchromosome linkers are found in cells, and that they are not an artifact of genome extraction.

We observed intact-cell CENP-B linkers for only about 8% of prometaphase CENP-B centromeric foci; it is possible that CENP-B containing linkers are relatively rare, and that there are a variety of different kinds of interchromosome linkers. rDNA-containing linkers represent one such class associated with rDNA-containing chromosomes (Potapova et al., 2019) (six in the human genome); possibly CENP-B-containing linkers are possessed by another subset of chromosomes in the human genome. Finally, the CENP-B-stained linkers observed in intact cells may be only parts of the linkers near the centromere itself, which are observable in the cells, but for isolated genomes, only represent a small fraction of stretched linkers.

In summary, we examined physical connections between mitotic chromosomes and found that chromosomes are mechanically tethered together by linkers associated with centromeric regions. Micromechanical study of the linkers shows a doubling force of ∼300 pN. Enzymatic treatment of interchromosome linkers held between two pipettes with applied tension demonstrates that the physical connection can be disrupted by DNase, but not by RNase, protease or spindle inhibitors, indicating the physical linker is connected by DNA elements. Immunofluorescence staining of proteins reveals Topo II is associated with the linkers in clusters spaced by approximately 0.4 Mbp. Experiments in intact cells confirm the existence of interchromosome linkers near the centromeres and suggest that the appearance of linkers in isolated genomes is not an artifact of their extraction from the cell (Fig. 7). We also observed that cell arrested during metaphase using microtubule-polymerization inhibitors lack interchromosomal linkers, possibly due to prolonged mitotic arrest causing DNA cleavage.

## Acknowledgements

This work was supported by the NIH by grants R35-GM161403 (JFM) and R16-GM153517 (LLH).

