## Supplemental Table and Figures for "Mitotic chromosomes are mechanically connected by chromatin-based interchromosome linkers"

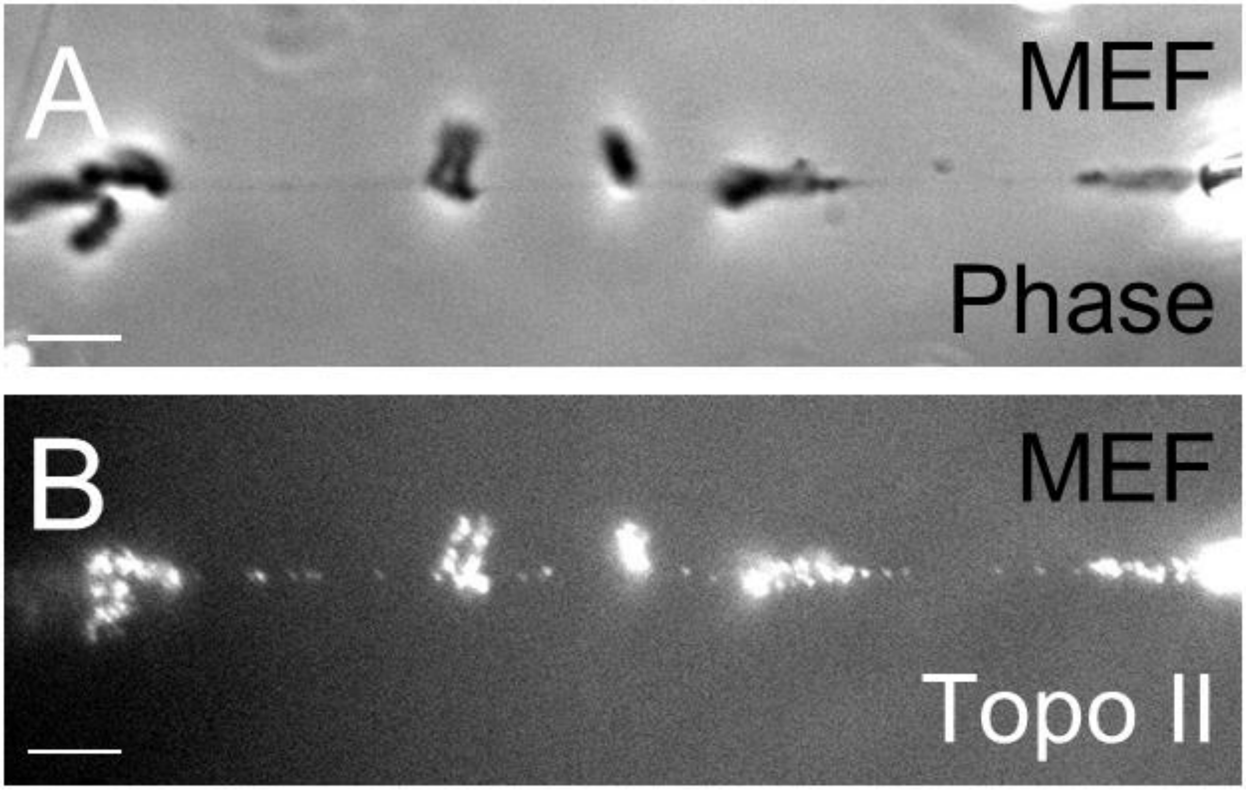


**Figure S1. Interchromosomal linkers in MEF cells are decorated by Topo II clusters.** Phase contrast (A) and immunofluorescence of Topo II (B) of the same chromosome clusters. Bar = 5 μm.

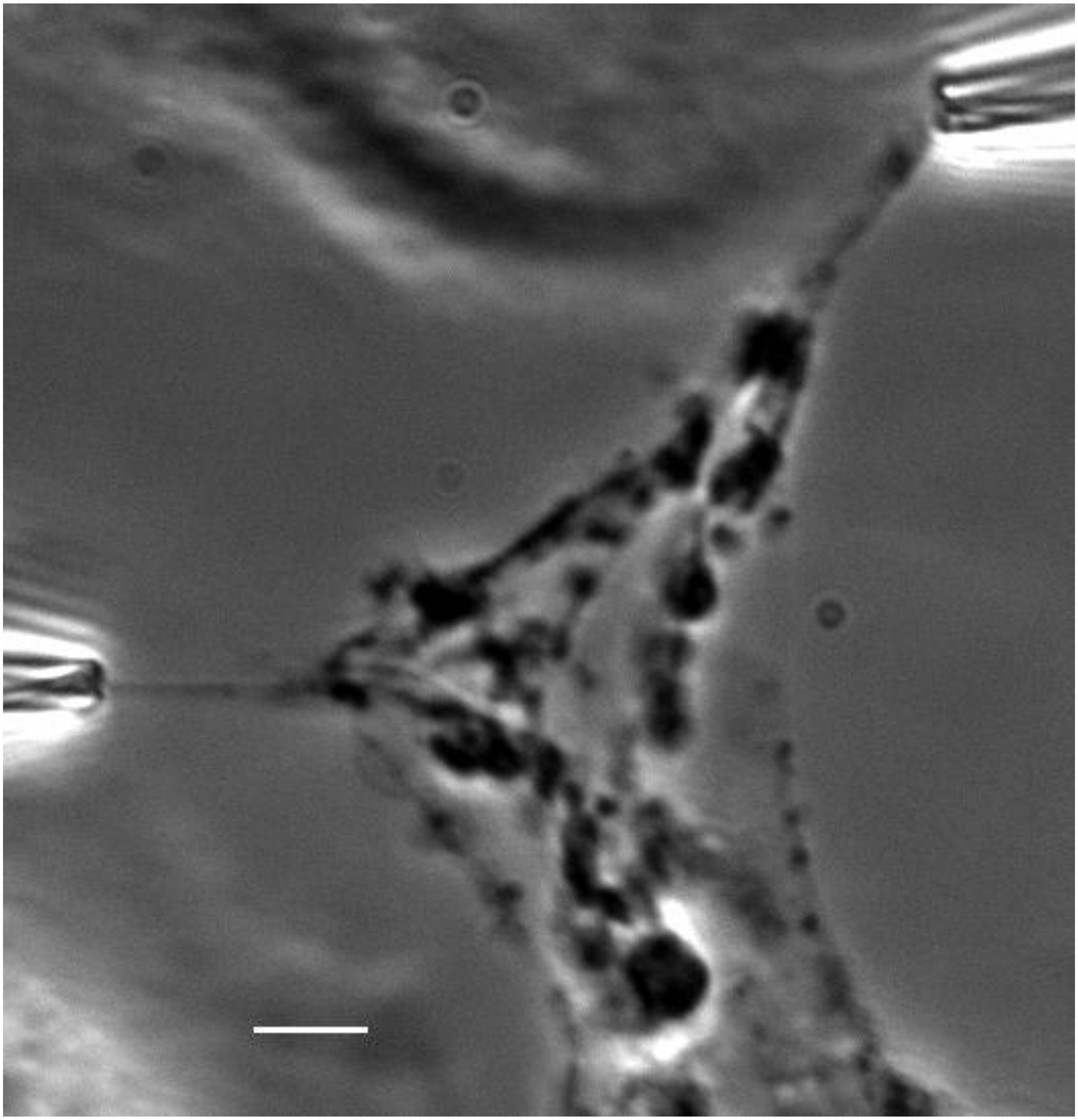

**Figure S2. Linkers occur in anaphase HeLa cells.** One half of a anaphase genome near one spindle pole (upper right) was pulled using glass micropipettes. Chromosomes are still connected by linkers as indicated by their alignment along the pulling direction (towards upper right), as in earlier stages of mitosis. Bar = 5 μm.

**Supplemental Table.**

Data for relaxed length, observed linker spring constants, and doubling forces for human HeLa mitotic interchromosome linkers. N=9. Cases * and ** correspond to Figs. 5C and 5D, respectively.

| Linker spring constant (pN/ μm) | Linker zero-force length (μm) | Linker doubling force (pN) |
|---|---|---|
| 35 | 5.3 | 188 |
| 52 | 12.6 | 650* |
| 85 | 2.9 | 250 |
| 33 | 4.0 | 130 |
| 78 | 2.2 | 175 |
| 30 | 8.0 | 240 |
| 24 | 6.7 | 160 |
| 94 | 4.3 | 405 |
| 50 | 3.2 | 160** |
